# DsbA-L protects against diabetic renal injury through the adipo-renal axis

**DOI:** 10.1101/2021.01.12.426410

**Authors:** Lingfeng Zeng, Ming Yang, Chun Hu, Li Zhao, Xianghui Chen, Yaping Wei, Huapeng Lin

## Abstract

Disulfide-bond A oxidoreductase-like protein (DsbA-L) is an adiponectin-interacting protein that is highly expressed in adipose tissue. The adipo-renal axis involves adipocyte release of signaling molecules that are recruited to kidney and regulate kidney function. We have found that the DsbA-L modulated the progression of diabetic nephropathy, but the precise mechanism of this modulation is unknown. Here, the transgenic mice overexpressing DsbA-L protein in fat (fDsbA-L) were used to verify that the renoprotective role of DsbA-L whether by adipo-renal axis. Mice were divided into four groups: a normal (Control) group, STZ induced diabetic mice, fDsbA-L mice and diabetic fDsbA-L mice (n=6). Diabetes was induced in mice by STZ 100mg/kg and continued HFD feeding for 12 weeks. Compared with the control group, the weight, blood glucose,and urine protein levels and the pathological changes in the kidney tissue of diabetic mice were increased significantly, accompanied by increased NLRP3,caspase-1, IL-1β, IL-18, FN, and Collagen1 mRNA and protein expression, which were reduced in diabetic fDsbA-L mice. Interestingly, the level of adiponectin in serum and kidney expression in diabetic mice was reduced significantly compared to that in the control group. However this change was reversed in diabetic fDsbA-L mice. These data suggest that the overexpression of DsbA-L in the adipocytes of mice can protect against diabetic renal injury through anti-inflammatory mediators,and may be mediated by the adipo-renal axis.

## 1. Introduction

Diabetic neprhropathy (DN) is one of the most serious microvascular complications in patients with diabetes mellitus, and is the primary cause of end-stage renal disease (ESRD), which is also one of the most leading cause of ESRD (ZengXiao and Sun 2019). DN is characterized by glomerular hypertrophy, mesangial expansion, K-W nodules, tubular and interstitial injury, and finally to the development of glomerulosclerois and renal fibrosis(Alicic Rooney and Tuttle 2017). However, the underlying mechanism of DN has not been fully elucidated. A variety of factors are involved in the pathogenesis of DN, including hemodynamic changes, genetic factors, oxidative stress, glucose metabolism disorders, cytokines and so on(Hu et al. 2015). However,after tightly controlling glucose and blood pressure in patients with DN, some patients still develop to ESRD. Recent studies have shown that inflammatory factors may be the important pathophysiological factors in the kidney injury in DN(Qiu and Tang 2016, Kim et al. 2016), but the precise mechanism is still not fully understand.

DsbA-L is a member of the glutathione S-transferase (GST) superfamily of antioxidant enzymes, which are involved in the primary cellular defense mechanism against reactive oxygen species(Hayes and Pulford 1995). Recent studies showed that DsbA-L is highly expressed in adipose tissue, and we have demonstrated the the downregulation of DsbA-L expression in the kidney tissues of DN in patients and in DN mice(Chen et al. 2019), the expression level of DsbA-L in mouse and human subjects is negatively correlated with proteinuria and podocyte and tubular damage in obesity and DN(Blackburn et al. 2011). These results indicate that the DsbA-L is associated with trhe renal injury of DN.

Furthermore, DsbA-L can affect the adiponectin secretion and polymerization as well as promote the expression of adiponectin and alleviate endoplasmic reticulum stress-induced adiponectin downregulation in adipose tissues, thus protecting against obesity and insulin resistance(Kadowaki et al. 2006). In addition, the adiponectin levels decreased in diabetic mice(Liu et al. 2012). Adiponectin is a secreted factor in adipose tissue that plays a critical role in anti-inflammatory, anti-fibrosis, and antioxidant processes and regulates glucolipid metabolism in DN(Yuan et al. 2014a). In addition, recent studies found that the adipo-renal axis may play an important role in kidney function, which means is that the adipocytes can release some signaling molecules such as adiponectin, leptin, IL-6 and TNF that are recruited to kidney tissues and mediate kidney pathophysiological changes(Zhu and Scherer 2018). However, whether DsbA-L in adipose tissue regulates the expression and release of adiponectin to protect diabetic kidney inflammatory injury needs to be further addressed.

## 2. Materials and methods

### 2.1 Animal experimental design

Transgenic mice overexpressing DsbA-L protein in fat (fDsbA-L) were donated by Professor Liu Feng from the University of Texas Health Science Center at San Antonio (UTHSCSA) (Transgenic Mice Core of UTHSCSA)(Liu et al. 2012). The genomic DNA of fDsbA-L mice was extracted by from the tail tissues to confirm gene overexpression, and it was used for identification of the genotype by PCR. The specific primers (upstream primer: ATCATTGCCAGGGAGAAC; downstream primer:TGCTTCAGGAGAGGAATC) were used for PCR amplification, followed by agarose gel electrophoresis and gel imaging. Experimental animals were divided into four groups: a normal (control) group, diabetic mice, fDsbA-L and diabetic fDsbA-L mice group (fDsbA-L+HFD+STZ)(n=6). Diabetic mice were fed a with high fat diet(HFD) for 4 weeks,followed by a single intraperitoneal injection of STZ (100mg/kg). After Three days after injection,fasting plasma glucose≥13.9mmol/L or random blood glucose≥16.7 mmol/L were taken as indicators of successful DN modeling, and then continue with HFD feeding was continued for 8 weeks. During the feeding period, blood glucose and body weight were measured every two weeks.At the end of the experiment, the blood and urine were collected and kidney tissue was harvested, and they was used for various experiments.

### 2.2 Serum and Urine biochemistry

Blood glucose levels were measured by a blood glucose monitor (Boehringer Mannheim, Germany). Urine albumin and serum adiponectin were detected by enzyme-linked immunosorbent assay (ELISA; Bethyl Laboratories, USA) following the manufacturer’s protocol.

### 2.3 Morphological studies

The renal tissue was isolated and fixed with 4% formalin, embedded in paraffin,and then cut into 4 μm thick sections. Hematoxylin and eosin (HE), periodic acid-Schiff (PAS), and Masson’s trichrome staining were performed on the paraffin sections as previously described (Yang et al. 2017).

### 2.4 Immunohistochemical Analysis

The related protein expression in the kidney tissues of mice was detected by Immunohistochemical (IHC) analysis. Four-micron-thick kidney sections were prepared, deparaffinized, rehydrated and then blocked with 3% H2O2. Using a microwave oven for antigen retrieval, the tissue sections were incubated with related antibodies against adiponectin(1: 100, ab22554, mouse monoclonal, Abcam, USA), caspase-1(1: 100,ab1872, rabbit polyclonal, Abcam, USA), IL-1β(1: 100,ab9722,rabbit polyclonal, Abcam, USA), and IL-18(1: 100,ab9722,rabbit polyclonal, Abcam, USA). DAB solution was then added to the renal tissues according to the manufacturer’s instructions.

### 2.5 Reverse transcription polymerase chain reaction

Total RNA from renal tissues was isolated using the Trizol kit following the manufacturer’s instructions. First-strand cDNAs were generated by two steps according to the reagent kit instructions. Then PCR amplification were performed. The protocol included a cycle at 95°C for 3 minutes; followed by 32 cycles of the following: 95°C for 45 seconds, 60°C for 45 seconds and 72°C for 60 seconds, and a final extension cycle at 72°C for 5 minutes(Yang et al. 2017). The PCR primers were as follows: FN (mouse,353bp) (sense, 5’-TTCCTGCACGTGTTTCGGAG −3’, antisense, 5’-TGTGCTGAAGCTGAGAACTAGG −3’);Collagen1 (mouse,285bp) (sense, 5’-CTTAAATGGAGGTGCAGGGCT −3’, antisense, 5’-CCCGCAGCCTTTTTGATAGC-3’); and β-actin (mouse,299bp) (sense, 5’-GACGGCCAGGTCATCACTAT-3’, antisense, 5’-CCACCGATCCACACAGAGTA-3’). Gel electrophoresis and gel imaging were then performed..

### 2.6 Western blot analysis

The protocol for the Western blot analyses was as follows: samples (20μg protein) were subjected to SDS–PAGE. When the proteins were transferred onto the nitrocellulose membranes, the blots were incubated with the specific antibodies. Autoradiograms were performed according to the manufacturer’s protocol.

### 2.7 Statistics

All statistical analyses were performed using SPSS 20.0 software. The results are expressed as the mean ± standard deviation (±SD) and were assessed by one-way analysis of variance (ANOVA). A value of p < 0.05 was considered as the significant difference between groups.

## 3. Results

### 3.1 Identification of the overexpression of disulfide-bond A oxidoreductase-like protein (DsbA-L) in mice and the levels of blood glucose, body weight,and urinary albumin

Transgenic mice of fDsbA-L were identified by PCR with specific primers. As shown in Figure1 A, a 324bp PCR product was observed, which indicated that exogenous DsbA-L was expressed in adipose tissue. Compared with the control group, the weight in diabetic mice was significantly increased, while body weight was reduced in diabetic fDsbA-L mice (Figure1 B). Similar results were also observed in the level of blood glucose (Figure1 C) and the degree of urinary albumin (Figure1 D). No significant difference in the weight of the mice, or the level of blood glucose was observed between the fDsbA-L group and the control group.

**Figure 1.**
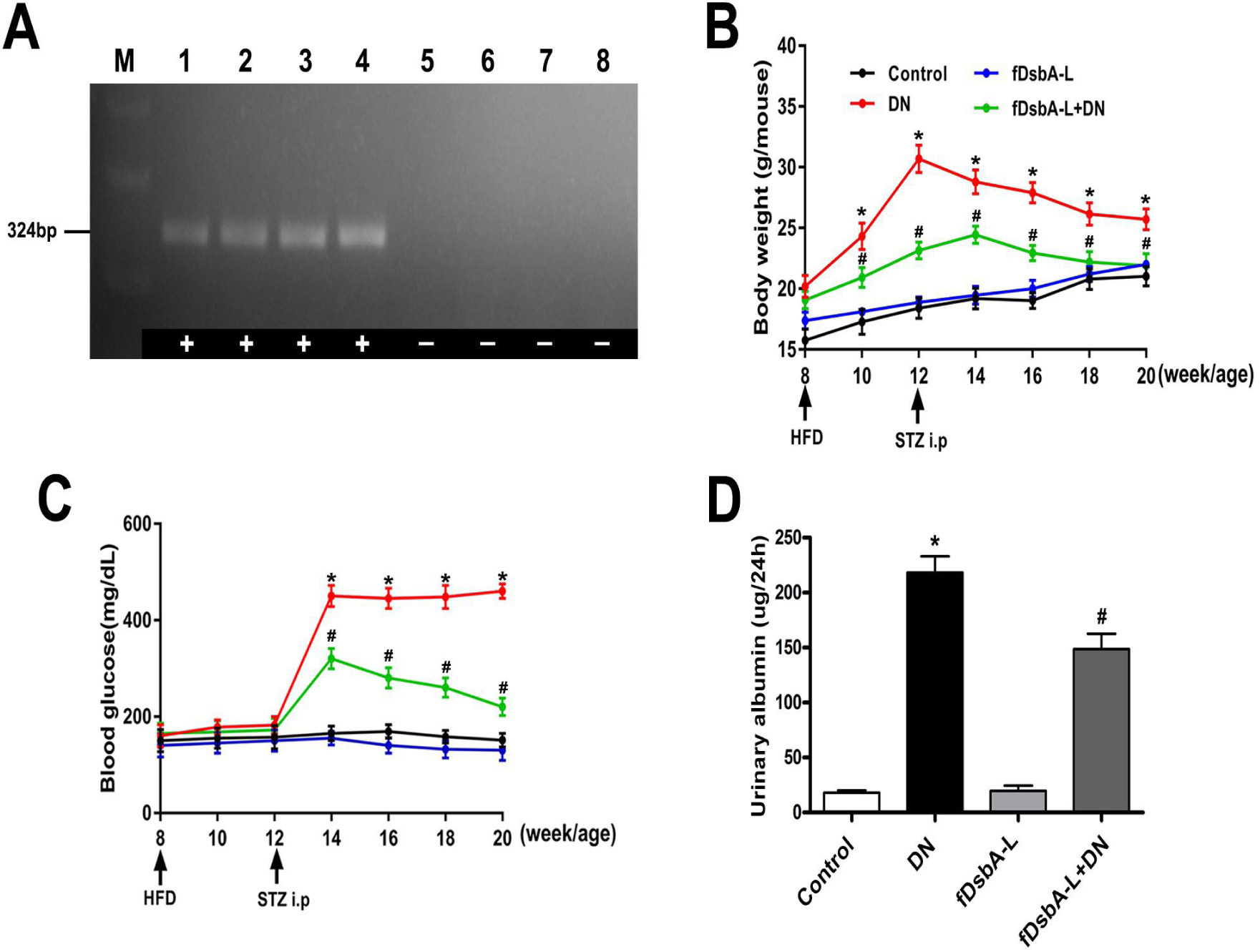
Genotyping fDsbA-L transgenic mice, and the assessment of body weight, blood glucose and urinary albumin. **A:** The gel showes that the polymerase chain reaction (PCR) amplification of tail genomic DNA from WT and fDsbA-L transgenic mice; **B:** Changes in body weight in mice; **C:**Changes in blood glucose in mice;**D**:Bar graph showing the levels of urinary albumin; (*p<0.05, compared with the control; #:p<0.05, compared with the diabetic mice.)

### 3.2 Expression of adiponectin and pathological changes in the kidneys of mice with fDsbA-L in adipose tissue

IHC staining showed that the expression of adiponectin was markedly decreased in the kidney tissue of diabetic mice compared with control mice, while the levels of adiponectin were elevated in diabetic fDsbA-L mice (Figure 2 A,B). In addition, a significantly decreased levels of serum adiponectin were found in diabetic mice compared with controls, while the level was enhanced in diabetic fDsbA-L mice(Figure 2 C);H&E staining showed that glomerular mesangial matrix proliferation was increased significantly in diabetic mice.However, the enhancement was alleviated in diabetic fDsbA-L mice (Figure 2 D a-d),PAS and Masson’s trichrome staining showed notably increased tubular brush border staining and tubulointerstitial fibrosis in the kidneys of diabetic mice when compared with control mice and reduced staining in diabetic fDsbA-L mice(Figure 2 D e-l). In addition, the glomerular injury and tubular damage scores were higher in diabetic mice than in control mice, and were reduced in diabetic fDsbA-L mice(Figure 2 E,F).These data suggested that the overexpression of DsbA-L in adipose tissue can reduce kidney injury, which may be related to up-regulating the expression of adiponectin.

**Figure 2.**
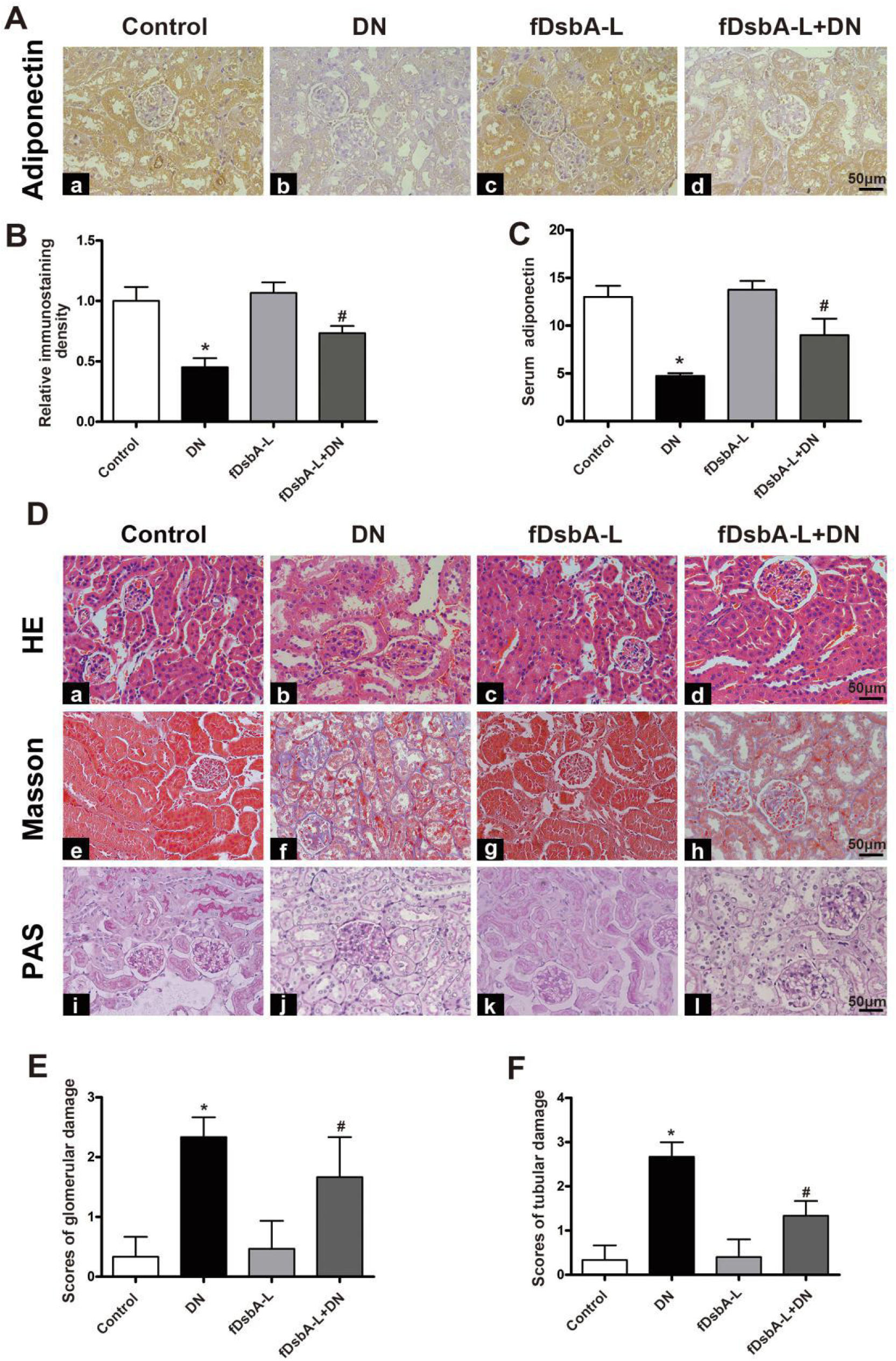
The expression of adiponectin in the kidney and renal morphological changes in diabetic mice. **A:**IHC staining showing that adiponectin expression in kidney tissue; **B:**Relative immunostaining density in the IHC staining analysis;**C:**The levels of adiponectin in the serum of different groups of mice as detected by ELISA. **D:** Kidney sections stained with H&E, PAS and Masson’trichrome; **E:** Bar graph representing the glomerular injury scores; **F:**Bar graph representing the tubular injury scores(*p<0.05, compared with the control group;#:p<0.05,compared with the diabetic group.)

### 3.3 Overexpression of DsbA-L in adipose tissue alleviated the levels of inflammatory cytokines in the kidneys of diabetic mice

IHC staining showed that NLRP3, caspase-1, IL-1β and IL-18 were mainly expressed in the cytoplasm of renal tubular epithelial cells. The inflammatory cytokines were notably increased in the kidneys of diabetic mice compared with the control group, whereas they were significantly decreased in the diabetic fDsbA-L group (Figure 3A,C). Western blot analysis revealed that the protein expression of NLRP3,caspase-1,IL-1β, IL-18 in the kidneys of the diabetic group was markedly elevated, while it was notably decreased in the diabetic fDsbA-L mice (Figure 3 D,E). These data indicated that the overexpression of DsbA-L in adipose tissue can alleviate inflammatory response in the kidneys of diabetic mice.

**Figure 3.**
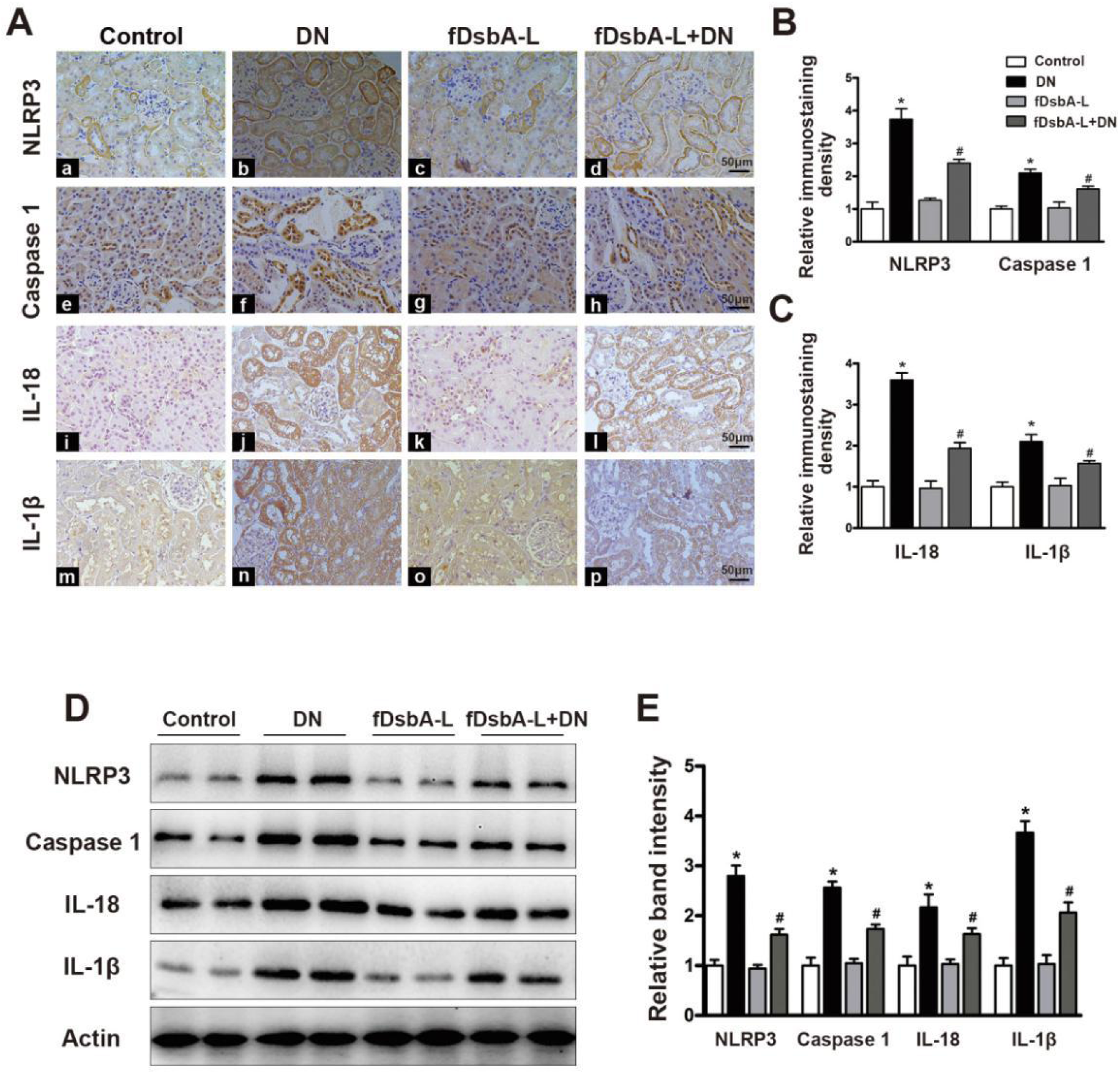
Effect of fDsbA-L on the expression of NLRP3,caspase-1,IL-1β and IL-18 in the kidneys of diabetic mice. **A:**IHC staining showed that the expression of NLRP3, caspase-1, IL-1β and IL-18 in the kidneys in the different groups of mice; **B-C:**Semiquantification of IHC staining for the expression of NLRP3, caspase-1,IL-1β and IL-18; **D:**Western blot analysis of the expression of NLRP3,caspase-1,IL-1β and IL-18 in the kidney tissue of mice from each group. **E**:The bar graph representing the relative band intensity by Western blot analysis. (*p<0.05, compared with the control group;#:p<0.05,compared with the diabetic group.)

### 3.4 Overexpression of DsbA-L in adipose tissue alleviated renal fibrosis in kidney tissue of diabetic mice

By real-time PCR, a significantly enhanced the expression of FN and Collagen1 mRNA was observed in the kidneys of diabetic mice compared with control mice.This observation was reversed in diabetic fDsbA-L mice(Figure 4A,B). IHC staining revealed that the protein expression of FN and Collagen1 were mainly expressed in the renal tubules and interstitium, and the pattern of FN and Collagen I expression was similar to the expression of inflammatory cytokines in the kidney (Figure 4C,D). These results were confirmed by Western blot analysis. The expression of FN and Collagen1 in the kidney was significantly decreased in diabetic fDsbA-L mice compared with diabetic mice (Figure 4E,F). These results suggest that the overexpression of fDsbA-L can alleviate renal fibrosis in the kidney tissue of diabetic mice.

**Figure 4.**
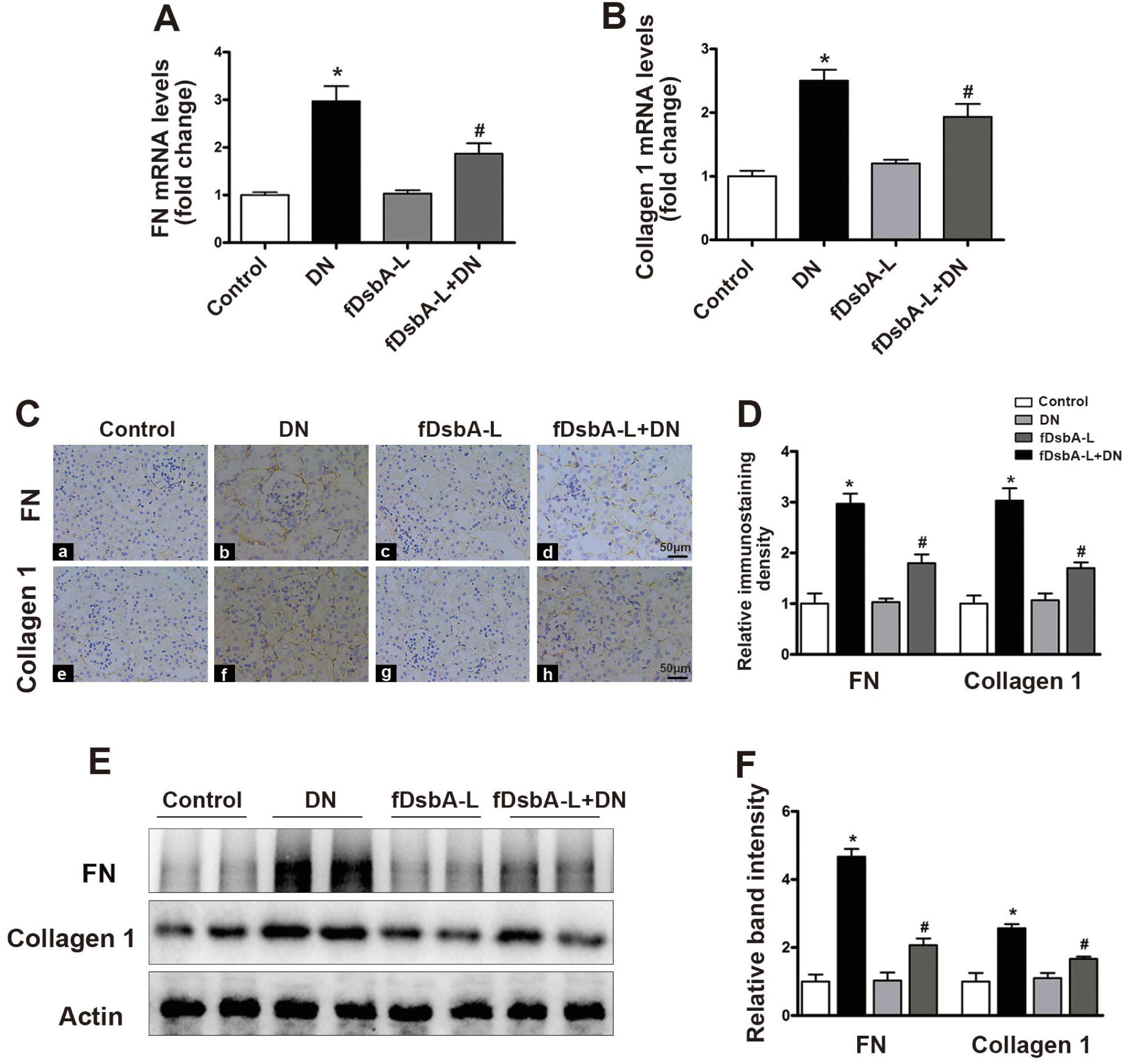
The expression of FN and Collagen I in the kidneys. **A-B:**The mRNA expression levels of FN and Collagen I in the kidneys in the different groups of mice as detected by real-time PCR; **C-D:** IHC experiment showing Collagen I,and FN expression in kidney sections from the different groups; **E:**Western blot analysis of the expression of FN and Collagen I in kidneys; **F:** Bar graph representing the relative band intensity by

## 4. Discussion

Diabetic nephropathy (DN) is a common kidney disease in people with diabetes, which is also a serious microvascular complication of diabetes and is the main cause of end-stage renal disease (ESRD) in developed and developing countries(RychlikMiltenberger-Miltenyi and Ritz 1998, Zeng et al. 2019). Over the last few decades, extensive efforts have been made to elucidate the pathogenesis of DN, and certain mechanisms have been identified, while others remains complex. Recent studies have shown that inflammatory factors may be the important pathophysiological factors in the kidney injury in DN (Matoba et al. 2019), but the precise mechanism is remains elusive.Recent studies indicates that the “adipo-renal axis” may play an important role in kidney function which means is that the adipocytes release some signaling molecules such as adiponectin, leptin, IL-6 and TNF that are recruited to kidney tissues and mediate kidney pathophysiological change(Zhu and Scherer 2018). In this study,we demonstrated for the first time that the overexpression of DsbA-L in the adipose tissue of mice has a renoprotective effect in diabetic mice via the “adipo-renal axis”.

DsbA-L, an adiponectin interactive protein recently identified by Dr.Liu (Liu et al. 2008), which localized in both the mitochondria and the ER in adipocytes and that its ER localization plays an important role in promoting adiponectin multimerization and stability in obesity-induced endoplasmic reticulum (ER) stress (Zhou et al. 2010).The overexpression of DsbA-L promoted adiponectin multimerization while the suppressing of DsbA-L expression significantly reduced adiponectin levels and secretion in 3T3-L1 adipocytes(Liu et al. 2012). In addition DsbA-L has been shown to have anti-inflammatory effects in a variety of diseases. Uno et al (Uno et al. 2013) have observed the protective effect of DsbA-L on renal damage induced by ROS in cynomolgus macaque.A study by our group has demonstrated that DsbA-L gene deficiency in mice aggravated lipid related renal injury in diabetic mice(Chen et al. 2019).

In addition, we also found that the knockout of the DsbA-L gene in mice have disturbed MAM integrity and modulated high-glucose-induced tubular cell damage(Yang et al. 2019). These results indicate that DsbA-L has a renoprotective role in diabetic mice. Since DsbA-L expression in adipose tissues which can regulate adiponectin and some preinflammatory expression and release, a recent study indicated that the adipo-renal axis plays an important role in regulating the pathophysiology role of the kidney.Thus, we proposed that DsbA-L in the adipocytes of mice has a renoprotective role in DN through the adipo-renal axis. As expected, we found that the overexpression of DsbA-L in adipocytes can reduce pathological changes which accompanied by increased adiponectin in serum and kidney tissue (Figure 1 and Figure 2).

The renoprotective role of DsbA-L whether through the regulation of the expression or release of adiponectin or inflammatory cytokines in adipose tissues is unknown. Adiponectin is a novel adipocyte-specific protein that plays a role in the development of insulin resistance and atherosclerosis(Qi et al. 2004, Mascarenhas-Melo et al. 2013). Adiponectin was also shown to be tightly related to the development of DN, Looker te al.(Looker et al. 2004)found that the level of adiponectin was decreased in the early stage of diabetic patients, Yuan et al.found that the intraperitoneal injection of exogenous adiponectin could reduce urinary albumin excretion and alleviate glomerular mesangial expansion(Yuan et al. 2014b, Yuan et al. 2010). It has been reported that DsbA-L expression,thereby increased adiponectin synthesis and secretion(Theodoratos et al. 2012).In addition, resveratrol, a powerful antioxidant that has enormous health benefits, for example,in diabetes and cardiovascular health, can upregulate the expression of DsbA-L, thereby upregulating the level of adiponectin in serum and adipose tissue(Wang et al. 2011).Here we found that the expression of serum and kidney tissue adiponenctin was signficantly decreased in diabetic mice compared to control mice, which was consistent with increased renal injury, which was rescued in mice with fat-specific= overexpression of DsbA-L (Figure 2). These data suggest that the overexpression of diabetic fDsbA-L has a renoprotective role in DN.

In this study, we found also that the expression of NLRP3, casepase1, IL-1β,and IL-18 was notably increased in diabetic mice,but reduced in diabetic fDsbA-L mice(Figure 3). A large number of studies have confirmed that inflammatory cytokines are important factors in diabetic kidney injury.It was found that the expression of NLRP3,IL-18and IL-1β was upregulated in the serum and kidney tissues of human and mice with DN,and the upregulation was correlated with kidney pathological injury and decreased renal function(Wan et al. 2019, Garrido et al. 2019). Skopinski et al.(Skopinski et al. 2005)found that the expression of IL-18 and IL-1β was significantly increased in the blood and urine of DN patients and was negatively correlated with the glomerular filtration rate and renal intersitial fibrosis. Interestingly, Guo et al(Guo et al.

2014)discovered that adiponectin retards the progression of DN by anti-inflammatory and anti-fibrosis actions in db/db mice. Importantly,Koshimura et al.(Koshimura et al. 2004) demostrated that the synthesis and secretion of adiponectin in adipose tissue and its early into the blood can alleviate diabetic microvascular complications.In contrast,the deletion of the gene for adiponectin would accelerate the renal inflammatory response and fibrosis in DN mice, which may occur through inhibiting the activation of the NF-kB pathway(Fang et al. 2015). In addition,the transplantation of adipose-derived mesenchymal stem cell sheets directly into the kidney suppressed the development of renal damage in DN rats (Takemura et al. 2019).Thus,we believe that increased DsbA-L expression in adipose tissues protects againsts diabetic kidney injury by increasing adiponectin secretion and reducing inflammatory cytokines release from adipose tissue, thereby increasing adiponectin levels and reducing inflammatory cytokine levels in renal tissue.

In summary, we demonstrated that the overexpression of DsbA-L in the adipose tissue of mice can protect against diabetic renal injury by increasing adiponectin secretion and reducing inflammatory cytokine release,which may mediated by the “adipo-renal axis”. Finally, the presented data should also provide momentum to intervene in the expression of DsbA-L in adipose tissue and provide a new therapeutic strategy for diabetic nephropathy treatment in the future.

## 6. Author contributions

F.L.Z. generated the data and performed the statistical analysis and drafted the manuscript. M.Y. and C.H. generated the data for the manuscript. and X.H.C. edited the manuscript and guided the statistical analysis and discussed the results of the manuscript. L.Z.and K.S.Y.conceived of the study, participated in its design, and coordination and wrote the manuscript. All authors read and approved the final manuscript.

## Cover letter

Dear Editors in Chief of Genetics,

We are submitting a manuscript entitled “DsbA-L protects against diabetic renal injury through the adipo-renal axis” by Zeng et al. for review and possible publication considerations in your journal.

Background: Diabetic nephropathy (DN) is one of the most serious microvascular complications in patients with diabetes mellitus, and is the primary cause of end-stage renal disease (ESRD). Recent studies have shown that inflammatory factors may play a critical role in the kidney injury of DN, but the precise mechanism is remains elusive. Recent studies have indicated that the “adipo-renal axis” may play an important role in the kidney through adipocyte release of molecules such as adiponectin, leptin, IL-6 and TNF, that are recruited to the kidney and mediate renal pathophysiological changes. Previous studies by our group and others have shown that DsbA-L has a renoprotective role in DN and have demonstrated that DsbA-L can affect adiponectin secretion and the expression of adiponectin. However, whether DsbA-L protects against diabetic kidney injury through the “adipo-renal axis” is unknown.

The Aim & Results: Transgenic mice overexpressing DsbA-L protein in fat (fDsbA-L)were used for this study to verify the renoprotective role of DsbA-L in the “adipo-renal axis”. The results showed that the pathological changes in the kidneys of diabetic mice were increased significantly compared to the control mice and were accompanied by increased NLRP3,caspase-1, IL-1b, IL-18, fibronectin(FN), and Collagen1 mRNA and protein expression,whereas the level of adiponectin in serum and kidney expression was reduced significantly. However, all of alterations were reduced in diabetic fDsbA-L mice. These data suggest that the overexpression of DsbA-L in the adipocytes of mice protects against diabetic renal injury by regulating of adiponectin expression and inflammatory cytokine secretion,which suggests that the “ adipo-renal axis “ may involved in the process.

We believe that this study will be of great interest to a broad audience in the diabetes mellitus and diabetic complications research field, including nephrologists and endocrinologists. The authors confirm that none of the material in the manuscript has been published or is under consideration for publication elsewhere.

I am the corresponding author and my address and other information as follows:

Thank you very much for your considerations.

Sincerely yours,

Huapeng Lin

Tel +86-023-63693626;

Fax +86-023-63693533;

